# Repetition of deliberate self-poisoning in rural Sri Lanka

**DOI:** 10.1101/344424

**Authors:** PHGJ Pushpakumara, SUB Thennakoon, TN Rajapakse, Ranil Abeysinghe, AH Dawson

## Abstract

Repetition of deliberate self harm is an important predictor of subsequent suicide. Repetition rates in Asian countries appear to be significantly lower than in western high income countries. The reason for these reported differences is not clear and has been suggested to due methodological differences or the impact of access to more lethal means of self harm. This prospective study determines the rates and demographic pattern of deliberate self-poisoning, suicide and fatal and non fatal repeated deliberate self-poisoning in rural Sri Lanka.

Details of deliberate self poisoning admission in all hospitals (n=46) and suicides reported to all the police stations (n=28) of a rural district were collected for 3 years, 2011-2013. Demographic details of the cohort of deliberate self-poisoning patients admitted to all hospitals in 2011 (N=4022), were screened to link with patient records and police reports of successive two years with high sensitivity using a computer program and manual matching was performed with higher specificity. Life time repetition was assessed in a randomly selected subset of DSP patients (n=438).

There were 15,914 DSP admissions and 1078 suicides during the study period. Within the study area the deliberate self poisoning and suicide population incidences were, 248.3/100,000 and 20.7/100,000 in 2012. Repetition rate for four weeks, one-year and two-years were 1.9% (95% CI 1.5-2.3%), 5.7% (95% CI 5.0 to 6.4) and 7.9% (95% CI 7.1 to 8.8) respectively. The median interval between two attempts were 92 (IQR 10 - 238) and 191 (IQR 29 - 419.5) days for the one and two-year repetition groups. The majority of patients used the same poison in the repeat attempt. Age and hospital stay of individuals with repetitive events were not significantly different from those who had no repetitive events. The two-year rate for suicide following DSP was 0.7% (95% CI 0.4-0.9%). Reported life time history of deliberate self harm attempts was 9.5% (95% CI 6.7-12.2%).

The low comparative repetition rates in rural Sri Lanka was not explained by higher rates of suicide or access to more lethal means or differences in methodology.

## Introduction

Deliberate self harm (DSH) is a major global public health problem. The World Health Organization (WHO) projects the worldwide yearly suicide mortality rate will increase to 1.53 million and it will be constitute 2.4 % of the total disease burden by 2020. (1). While there is significant variation of suicide rates between countries Sri Lanka’s suicide rates have remained amongst the highest in the world, (2, 3).

A recent meta-analysis estimated that one in 25 patients presenting to hospital for self-harm will suicide in the next 5 years. (4).Understanding factors that influence the rate and pattern of repetition of self harm has the potential to inform prevention strategies and optimal follow-up after a self-harm episode. There appears to be geographic differences in the 1 year non-fatal repetition rates. In European studies 1 year non-fatal repetition rates was estimated as 17.1% (95% CI 15.9-18.4) while it was lower in Asia (10.0%, 95% CI 7.3-13.6). (4). Possible proffered reasons for this included methodological weakness of the Asian studies, higher lethality of self-poisoning and longer hospital stay(4). It was suggested that identifying the reasons for this variation could provide insights into optimal configuration of health care services (4).

This prospective study determines the four weeks, one year and two year rates of fatal and non-fatal repeated self harm and estimates life-time repetition rate and pattern in deliberate self poisoning in the Kurunegala District (KD), of Sri Lanka, which was conducted as a part of comprehensive analysis of DSH and suicide in KD (5).

## Methods

### Study Setting and Design

This study was conducted in the predominately rural agricultural KD in Sri Lanka. The district has a population of 1.6 million(6) who have free access to 46 government hospitals; 45 District Hospitals and the tertiary Teaching Hospital Kurunegala (THK)(7). Both in-hospital and community deaths from suicides from any causes are reported to district police stations (n=28)

A prospective cohort of all hospital presentations following deliberate self-poisoning (DSP) to government hospitals within the KD was established between 1^st^ January 2011 to 31^st^ December, 2013 as part of a study of use of treatment guidelines (Sri Lanka Clinical Trial Registry No. SLCTR/2010/ 008). This study on self-harm repetition utilized this cohort for hospital data and in addition collected data on all suicides reported to district police stations, to identify fatal repetitions. A randomly selected subset of patients and their bystanders were interviewed to determine self-reported lifetime repetition rate and pattern.

### Study Recruitment Prospective Repetition Cohort

Identification, demographic and clinical details of all DSP admissions were collected for the study. In THK all patients were enrolled into the cohort at the time of admission, by fulltime study doctors employed as clinical research assistants. Patients were seen at least daily until discharge or death. At the other 45 district hospitals data was extracted from patient medical record by tertiary postgraduate research assistants and entered into a study data base along with a scanned copy of the medical record of the patient’s admission to facilitate audit. Hospitals were visited every 2-4 weeks depending upon the size of the hospital. Typically, all relevant admission records had been left aside at each hospital to facilitate case finding but in each hospital the admission ledgers were also reviewed to ensure all relevant medical records had been identified.

Details of all suicides reported to police stations were collected by visiting all 28 police stations in the district. Data was retrieved from suicide registers at each police station, by tertiary educated postgraduate research assistants, for the same period.

Within the cohort, the patients index admission was their first admission to any study hospital between 1^st^ January 2011 to 31^st^ December, 2011. Following the index admission, the study database was interrogated for repeat presentations for a period of two years to hospitals or police stations. As there is no unique patient medical record number within the provincial health system, identification of inter-hospital transfers and repeat presentations required individual identity linkage.

Initial linkage utilized surname, at least one of the other names, sex and age as mandatory fields and residential address as an optional field, for confirmation of the matching. A five step method, which has been adapted from a English-Sinhala transliteration system (8) and a process of matching names in Sinhala (9), was used in screening to generate possible spelling combinations of surname, other names and village/address; transliteration into Sinhala, decomposing Sinhala words, single and multi character replacements, generation of possible spelling combinations of Sinhala words by combining replaced characters and transliteration into English. Semi-automated stepwise data matching and filtering process was followed for record linking. Records were screened for links with high sensitivity using the computer, then subsequent manual confirmation of any screened results

### Study Recruitment Lifetime Recalled Repetition Cohort

Lifetime recalled previous self-harm was conducted in a randomly selected cohort of patients admitted to THK following DSP. Patients were randomly selected using a computer program from blocks of 7 consecutively admitted consenting DSP patients within a consecutive eighteen months from 1^st^ July 2011. Immediately prior to discharge patients received a structured interview by medical graduates research assistants who were l trained for the data collection. Patients were asked to recall previous episodes of self-harm (e.g. poisoning, hanging, and drowning). Interview information was verified through a close relative or someone well aware about the patient.

### Data analysis

Data were entered onto a Microsoft Access database and analyzed using SPSS version 23.

### Ethics statement

Ethics approval was obtained from Ethics Review Committee, Faculty of Medicine, University of Peradeniya for ‘A clustered RCT of educational interventions on treatment of patients with acute poisoning in rural Asian hospitals’. Ethical approval for additional data collection was obtained from the Ethics Review Committee, Faculty of Medicine and Allied Sciences, Rajarata University of Sri Lanka. The study was conducted with the support of the Provincial Department of Health Care and nutrition, of the North Western Province and Department of Police, Sri Lanka.

## Results

### Prospective Cohort Study

A total of 15,914 and 1,078 records were collected from hospitals and police stations respectively, for the 3 years. THK received 53% of all DSP cases of the district either as direct admission or following transfer. Follow-up cohort consisted of 4,022(50.8% males and 49.2% females) patients after removing double counting due to inter hospital transfers, with a median age of 23 years. The highest proportion of the cohort, 35.5% (95% CI 34.7 to 36.2), presented following ingestion of agro-chemicals. This was followed by overdose of medications, 32.9% (95% CI 32.1 to 33.6), and ingestion of oleander seeds, 15.2% (95% CI 14.6 to 15.7). In 2012 the DSP incidence in KD was 248.3/100,000 (95% CI 240.6 to 255.9) and male:female ratio was 1.1 (Table 1).

**Table 1:** Age standardized DSP incidences in 2012 in Kurnegala District among males and females, with male to female ratio.

|  | Age adjusted DSP Incidence in 2012 per 100,000 Population*<br>(95% CI) |  |  | Male:Female<br>Ratio |
| --- | --- | --- | --- | --- |
|  | Male | Female | Total |  |
| 10-14 | 80.7<br>(58.3 to 103.1) | 188.9<br>(154.3 to 223.4) | 134.3<br>(113.8 to 154.8) | 0.4 |
| 15 - 19 | 634.4<br>(570.9 to 697.8) | 1304.0<br>(1213.6 to 1394.3) | 971.4<br>(916.1 to 1026.7) | 0.5 |
| 20 - 24 | 769.6<br>(694.3 to 844.9) | 760.2<br>(688.0 to 832.4) | 764.7<br>(712.6 to 816.9) | 1.0 |
| 25 - 29 | 414.4<br>(360.7 to 468.1) | 371.3<br>(323.6 to 418.9) | 391.5<br>(355.8 to 427.2) | 1.1 |
| 30 - 34 | 317.2<br>(272.6 to 361.7) | 231.5<br>(195.2 to 267.9) | 272.4<br>(243.9 to 300.9) | 1.4 |
| 35 - 39 | 292.2<br>(247.3 to 337.0) | 144.0<br>(113.4 to 174.6) | 216.0<br>(189.1 to 242.9) | 2.0 |
| 40 - 44 | 243.3<br>(201.8 to 284.8) | 88.8<br>(64.4 to 113.1) | 163.8<br>(140.1 to 187.6) | 2.7 |
| 45 - 49 | 231.2<br>(189.9 to 272.6) | 56.7<br>(37.1 to 76.4) | 140.4<br>(118.0 to 162.7) | 4.1 |
| 50 - 54 | 175.4<br>(138.7 to 212.0) | 54.2<br>(34.8 to 73.6) | 111.8<br>(91.6 to 132.0) | 3.2 |
| 55 - 59 | 146.2<br>(110.6 to 181.7) | 23.5<br>(10.2 to 36.8) | 80.6<br>(62.6 to 98.6) | 6.2 |
| 60 - 64 | 118.0<br>(83.1 to 152.9) | 36.2<br>(18.4 to 53.9) | 73.6<br>(55.0 to 92.2) | 3.3 |
| 65 - 69 | 125.6<br>(78.2 to 172.9) | 36.3<br>(13.8 to 58.8) | 75.5<br>(51.1 to 99.8) | 3.5 |
| 70 - 74 | 128.6<br>(70.8 to 186.4) | 15.3<br>(-2.0 to 32.7) | 64.0<br>(37.3 to 90.8) | 8.4 |
| 75 - 79 | 120.4<br>(49.3 to 191.6) | 14.2<br>(-5.5 to 34.0) | 56.1<br>(25.6 to 86.6) | 8.5 |
| 80 & over | 188.8<br>(99.0 to 278.6) | 37.1<br>(4.6 to 69.6) | 97.8<br>(56.9 to 138.7) | 5.1 |
| Overall | 257.7<br>(246.4 to 269.0) | 239.5<br>(229.1 to 249.9) | 248.3<br>(240.6 to 255.9) | 1.1 |

\*DSP incidences were calculated based on the 2012 DSP events

A total of 77 (n=44, 57% were males) had a repeat self-harm event within the first four weeks from the indexed event. Repetition rate for the four week period was 1.9% (95% CI 1.5-2.3%). The median age of those who repeated within the first four weeks was 27 years (IQR 19 – 44 years).

There were 179 (56.3%) males and 139 (43.7%) females with repetition of self-harm within two years. One-sample binominal test showed that repetitive events were significantly common among males (p=0.03) and being a male carried a 1.3 fold excess risk for repetitive attempts (OR 1.3, 95% CI 1 to 1.6).A majority, 290 (91.2%), had only one repetitive attempt, 24 (7.5%) had two, 3 (0.9%) had three and one (0.3%) had four during that period. One year and two year repetition rates were 5.7% (95% CI 5.0 to 6.4) and 7.9% (95% CI 7.1 to 8.8) respectively. The median age of males who repeated self-harm within the two year follow-up period was 28 years (IQR 20 – 40 years) and for females it was 19 years (IQR 16 – 25 years). For more than one fifth (22.3%) of males and nearly half (48.9%) of the females repetition occurred in the 15-19 year age group, which is an over representation compared to the cohort, male 22% and female 37%. Mann Whitney test analysis showed that compared to the ages of males, females were younger, p < 0.0001. One sample chi-square test showed that probabilities of having a repetitive event among age categories were significantly different, p < 0.0001. Relatively higher repetition rates were reported among younger age groups in females and opposite pattern in males (Table 2, Figs 1 and 2). T-test analysis showed that the ages of individuals who had repetitive events and who had no repetitive events, do not different significantly, p= 0.33.

**Fig 1.**
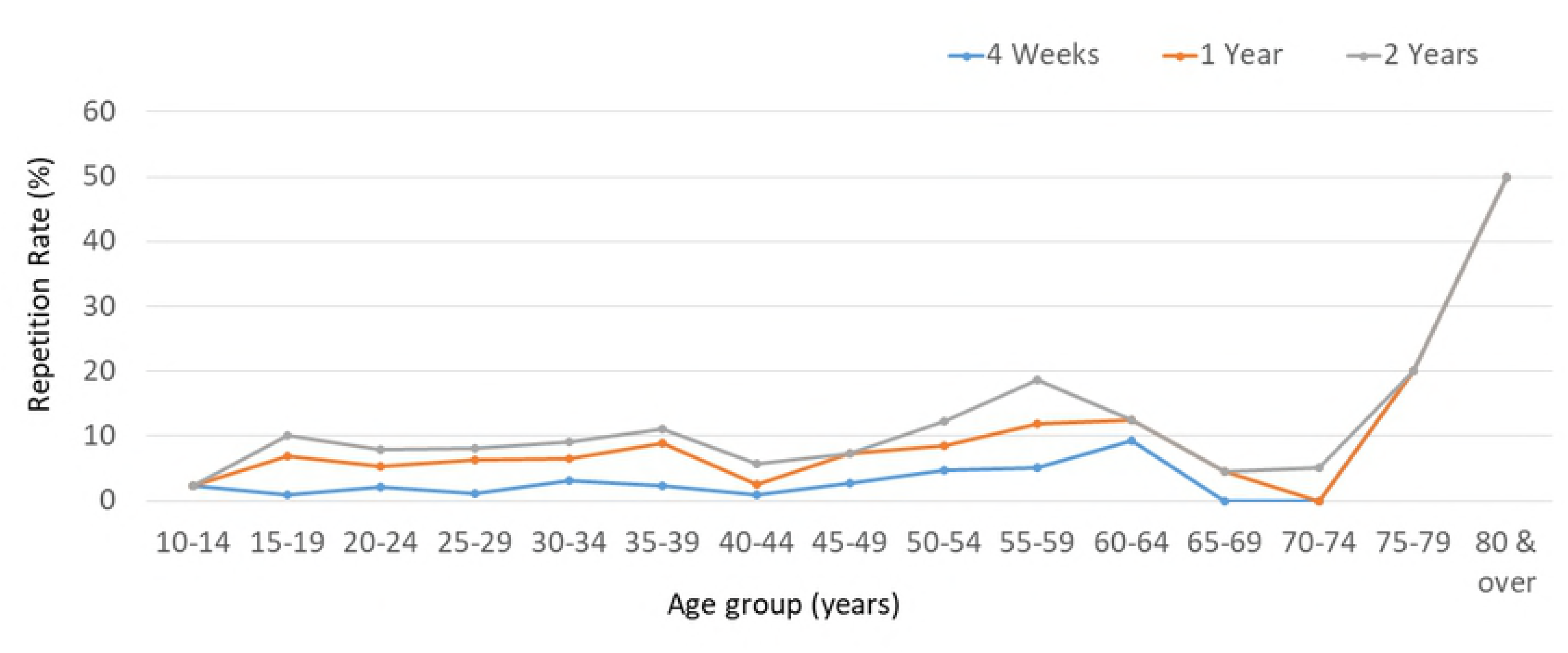
shows the four weeks, one year and two year repetition rate of males by age group and sex.

**Fig 2.**
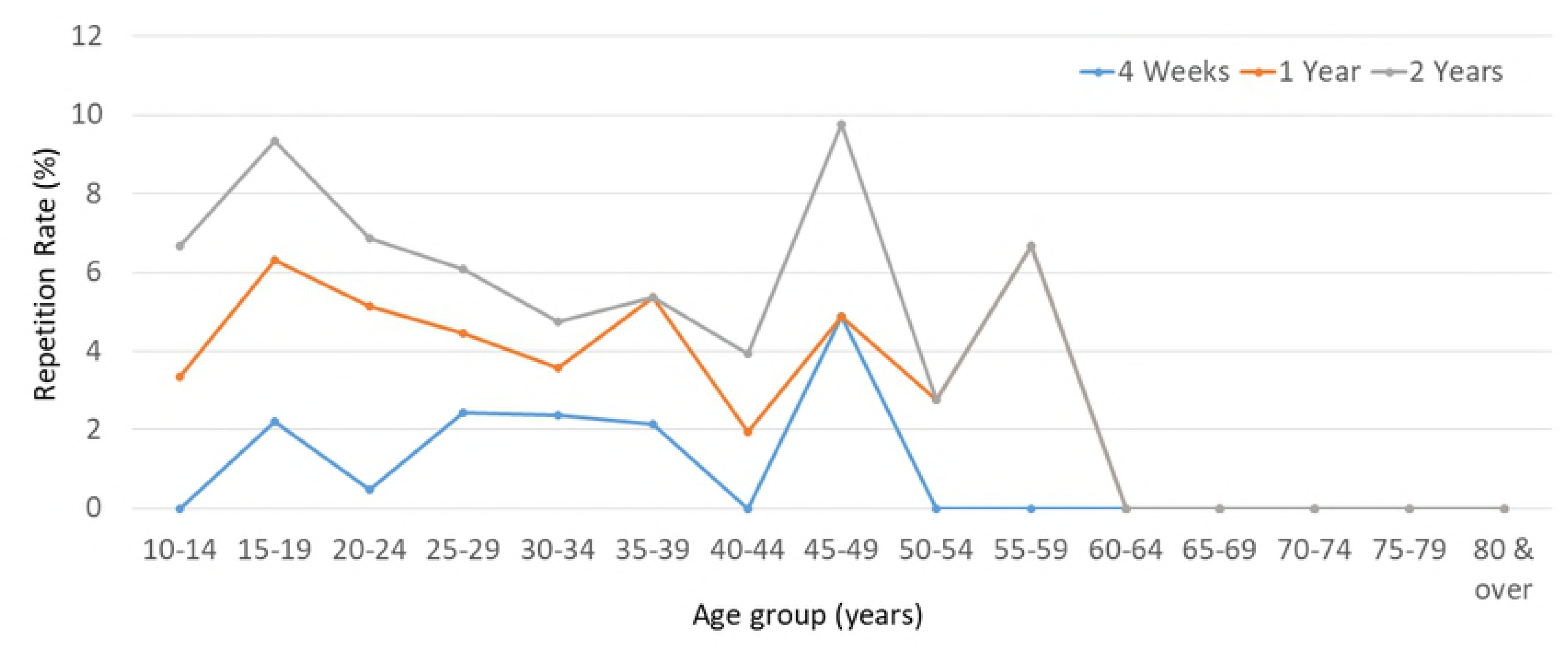
shows the four weeks, one year and two year repetition rate of females by age group and sex.

The average intervals between two consecutive events were 246.8 days (SD 223.4) among males and 238.5 days (SD 207.0) among females, and this difference was not significant, p=0.7. One-sample Kolmogorov-smirnov test showed that intervals between index event and the first repetitive event were not normally distributed, p < 0.0001. Fig 3 shows the cumulative probability of the first repetitive DSP events in first two years by sex. Median times to repetition within 1 year and 2 years were, 92 (IQR 10 - 238) and 191 (IQR 29 - 419.5) days. The highest risk for repetition was observed in the initial one week period, where 17% (n=54) repetitive attempts occurred. 9.1% (n=29) had re-attempted on the following day of the indexed event.

**Fig 3.**
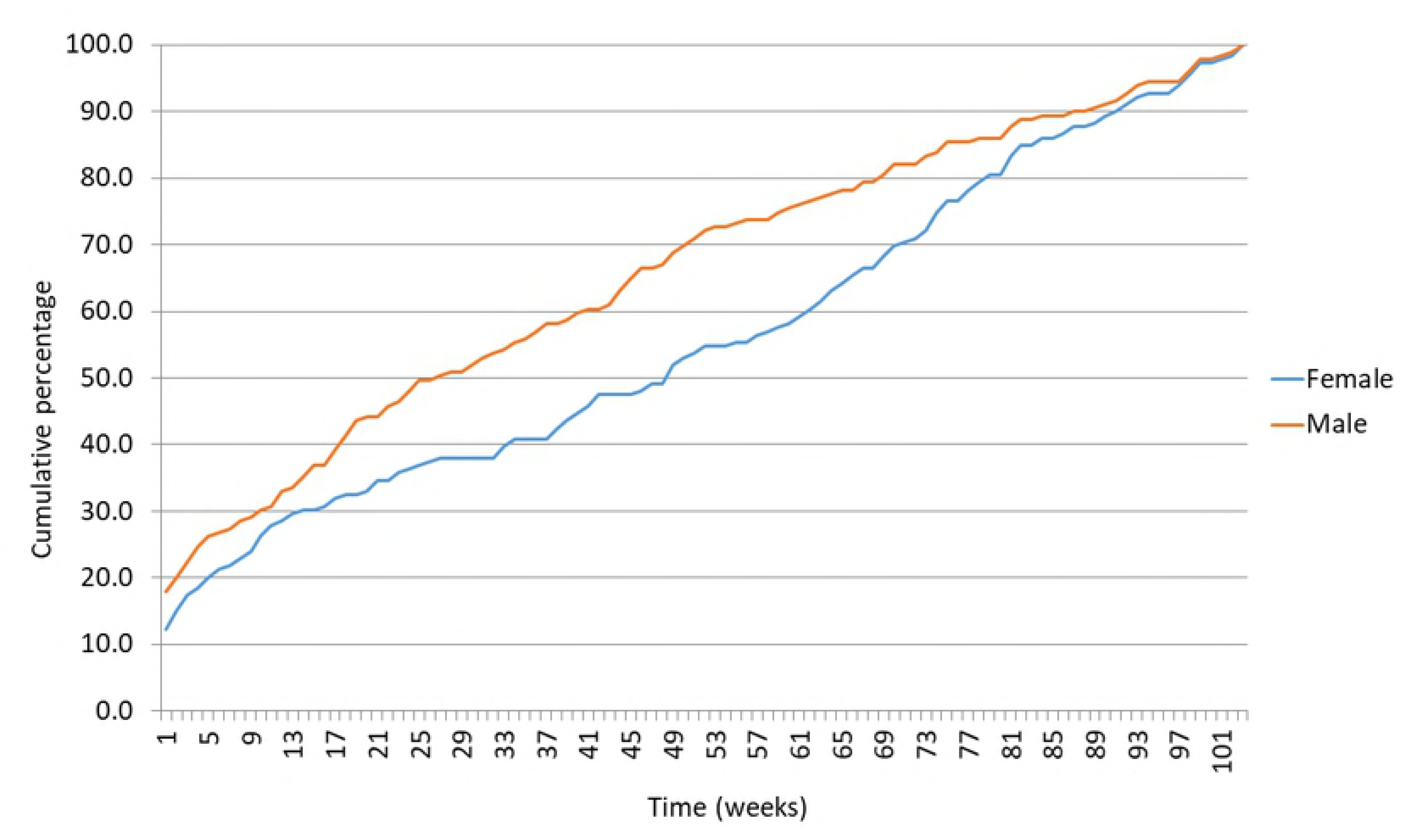
shows the cumulative probability of the first repetitive DSP events during their first two years, by sex.

The first repetitive event was a fatal for 28 (8.8%) individuals. None of the second, third or fourth repetitive events were fatal. The two year rate for suicide following DSH was 0.7% (95% CI 0.4-0.9%). All fatal repetitive events were reported among males. The mean age for those who carried out fatal repetitive events was 49.7 (SD 15.3) and those who died were significantly older compared to those who had non-fatal outcome, p<0.0001. Nearly 40% of fatal two year repetitive events occurred within the first week and 50% within first 3 weeks following the indexed event.

The pattern of type of poison used for the repetitive events was similar to the pattern of the cohort. 60% of individuals who ingested agro-chemicals used the same method for the next consecutive event. Nearly half (47%) and more than half (55%) of individuals who overdosed medicines and ingested oleander seeds used the same method for the next consecutive event. A majority, 24 (85.7%), of fatal repetitions were due to poisoning; two due to oleander and 22 agro-chemicals. One (3.6%) was due to hanging and 3 (10.7%) were not classified. More than three forth (77%) of individuals who ingested agro-chemicals at the fatal repetitive event used the same method at the index event. Table: 3 summarizes the findings.

**Table 3:** Fatal and non-fatal repetitive events by the type of the poison and pattern of use at the subsequent events.

| Type of poison ingested | Individuals in Cohort | Individuals with Repetitive events |  |  |  |
| --- | --- | --- | --- | --- | --- |
|  |  | Use at the indexed Event | Same Method used in any two consecutive events | Used for the fatal events | Same Method used in the fatal and Indexed event |
|  | % (95% CI) | n (%) | n (%) | n (%) | n (%) |
| Agro-Chemical | 35.5 (34.7 to 36.2) | 127 (39.9) | 76 (48.7) | 22 (91.7) | 17 (94.4) |
| Medicine | 32.9 (32.1 to 33.6) | 93 (29.2) | 44 (28.2) | 0(0) | 0(0) |
| Oleander | 15.2 (14.6 to 15.7) | 60 (18.9) | 33 (21.1) | 2 (8.3) | 1 (5.6) |
| Hydrocarbon | 4.6 ( 4.3 to 4.9) | 10 (3.1) | 2 (1.3) | 0 (0) | 0 (0) |
| Alkali | 0.1 ( 0.1 to 0.2) | 1 (0.3) | 0 (0) | 0 (0) | 0 (0) |
| Rodenticide | 0.8 (0.7 to 1.0) | 3 (0.9) | 0 (0) | 0 (0) | 0 (0) |
| Fertilizers | 0.6 (0.5 to 0.7) | 1 (0.3) | 0 (0) | 0 (0) | 0 (0) |
| Acid | 0.2 (0.1 to 0.3) | 0(0) | 0 (0) | 0 (0) | 0 (0) |
| Other Chemical | 0.9 (0.7 to 1.0) | 3 (0.9) | 0 (0) | 0 (0) | 0 (0) |
| Oleander & Medicine | 0.1 (0.1 to 0.2) | 1 (0.3) | 0 (0) | 0 (0) | 0 (0) |
| Medicine & Agro-Che. | 0.3 (0.2 to 0.4) | 0 (0) | 0 (0) | 0 (0) | 0 (0) |
| Combined Agro-Che. | 0.07 (0.03 to 0.11) | 0 (0) | 0 (0) | 0 (0) | 0 (0) |
| Other Combination | 0.5 (0.4 to 0.6) | 0 (0) | 0 (0) | 0 (0) | 0 (0) |
| Unknown | 8.2 (7.8 to 8.6) | 19 (6) | 0 (0) | 0 (0) | 0 (0) |
| Total | 100 | 318 (100) | 155 (100) | 24 (100) | 18(100) |

The median hospital stay of DSP patients managed at peripheral hospitals, for both who had and did not have repetitive attempts, were two days (Table 4). The duration of the hospital stay did not differ significantly depending on the type the poison. Further, it did not show a significant association with the occurrence of repetition of self-harm. It showed that 4.1% (95% CI: 3.2 – 5.2%) and 2.9 % (95% CI: 2.1-3.7%) of patients admitted to peripheral hospitals and THK were discharged at the same day. 23.6% (95% CI: 21.6 – 25.7%) and 17.2% (95% CI: 15.5 – 19.0%) of patients were discharged the following day respectively.

**Table 4:**
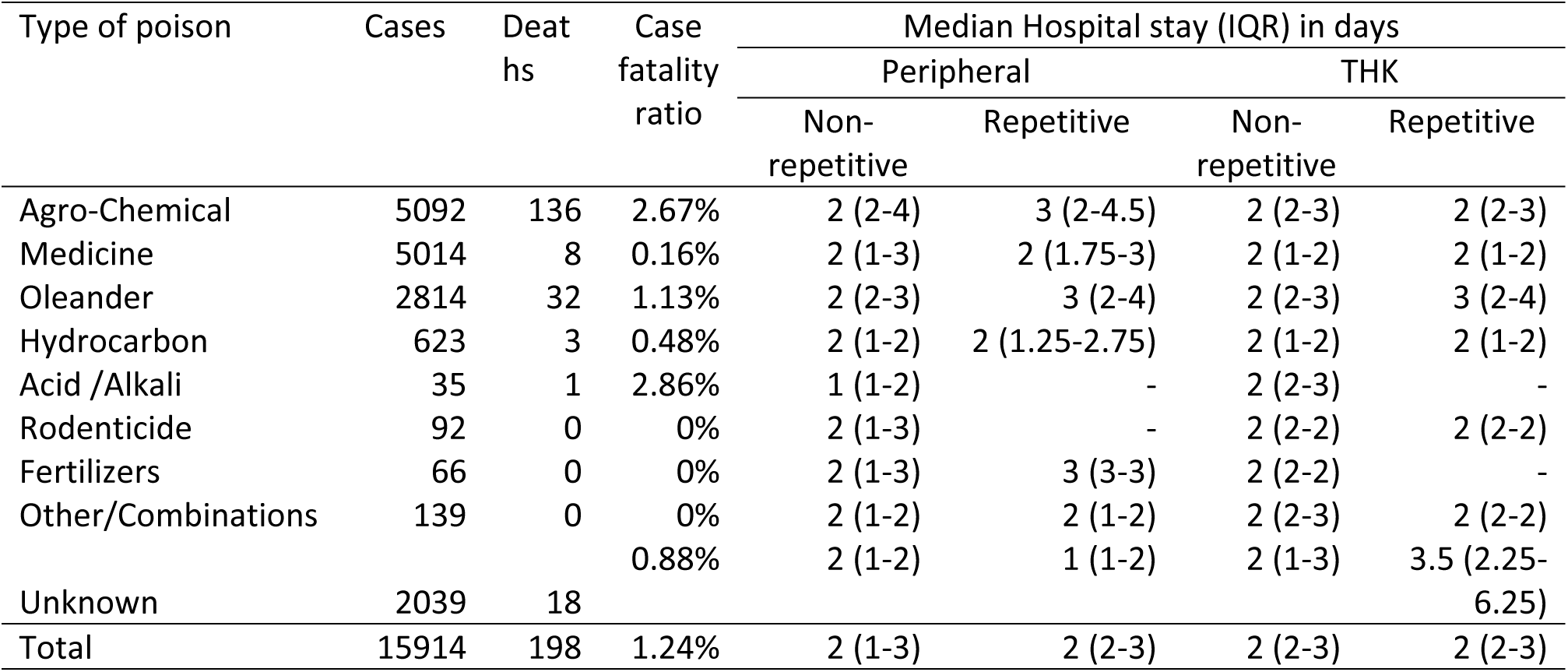
Duration of hospital stay and case-fatality ratio by type of poison

There were 1,078 suicides in the district by all methods in 2011 to 2013 (Table 5). It showed that only 31.2% of male and 33.3% of female suicides by poisoning reported to hospitals. Suicide incidence in KD was 20.7/100,000 (95% CI 18.5 to 22.9) and the male:female ratio was 4.4 (Table 6).

**Table 5:** Suicides in KD by method and sex

| Methods | Male |  | Female |  | Total |  |
| --- | --- | --- | --- | --- | --- | --- |
|  | N | % | N | % | N | % |
| Burning | 5 | 0.6 | 7 | 3.9 | 12 | 1.1 |
| Stabbing/Cutting with a sharp weapon | 2 | 0.2 | 1 | 0.6 | 3 | 0.3 |
| Drowning | 24 | 2.7 | 18 | 10.1 | 42 | 3.9 |
| Gun shot | 2 | 0.2 | 0 | 0.0 | 2 | 0.2 |
| Hanging | 269 | 29.9 | 30 | 16.9 | 299 | 27.7 |
| Jump to motor vehicle | 1 | 0.1 | 0 | 0.0 | 1 | 0.1 |
| Jump to Train | 43 | 4.8 | 7 | 3.9 | 50 | 4.6 |
| Oleander | 35 | 3.9 | 27 | 15.2 | 62 | 5.8 |
| Pesticide | 499 | 55.4 | 84 | 47.2 | 583 | 54.1 |
| Pesticide & Drowning | 1 | 0.1 | 0 | 0.0 | 1 | 0.1 |
| Other | 11 | 1.2 | 4 | 2.2 | 15 | 1.4 |
| Not Recorded | 8 | 0.9 | 0 | 0.0 | 8 | 0.7 |
| Total | 900 | 100.0 | 178 | 100.0 | 1078 | 100.0 |

**Table 6:** Age standardized suicide incidences in KD in 2012 among males and females

|  | Age adjusted Suicide Incidence per 100,000 population* (95% CI) |  |  | Male:Female<br>Ratio |
| --- | --- | --- | --- | --- |
|  | Male | Female | Total |  |
| 10-14 | 0.0 | 4.9 (-0.6 to 10.5) | 2.4 (-0.3 to 5.2) | - |
| 15 - 19 | 13.2 (4.1 to 22.4) | 14.7 (5.1 to 24.2) | 13.9 (7.3 to 20.6) | 0.9 |
| 20 - 24 | 40.3(23.1 to 57.5) | 19.6 (8.0 to 31.2) | 29.6 (19.3 to 39.8) | 2.1 |
| 25 - 29 | 25.3 (12.1 to 38.6) | 1.6 (-1.5 to 4.7) | 12.7 (6.3 to 19.1) | 15.9 |
| 30 - 34 | 26.0 (13.3 to 38.8) | 17.8 (7.7 to 27.9) | 21.7 (13.7 to 29.8) | 1.5 |
| 35 - 39 | 25.1 (11.9 to 38.2) | 6.8 (0.1 to 13.4) | 15.7 (8.4 to 22.9) | 3.7 |
| 40 - 44 | 47.9 (29.5 to 66.3) | 5.2 (-0.7 to 11.1) | 26.0 (16.5 to 35.4) | 9.2 |
| 45 - 49 | 67.4 (45.1 to 89.8) | 8.9 (1.1 to 16.6) | 36.9 (25.5 to 48.4) | 7.6 |
| 50 - 54 | 77.7 (53.3 to 102.1) | 3.6 (-1.4 to 8.6) | 38.9 (27.0 to 50.7) | 21.5 |
| 55 - 59 | 45.0 (25.3 to 64.7) | 0.0 | 20.9 (11.8 to 30.1) | - |
| 60 - 64 | 75.1 (47.3 to 102.9) | 11.3 (1.4 to 21.2) | 40.5 (26.7 to 54.3) | 6.6 |
| 65 - 69 | 74.4 (37.9 to 110.9) | 10.9 (-1.4 to 23.2) | 38.8 (21.3 to 56.2) | 6.8 |
| 70 - 74 | 108.3 (55.2 to 161.3) | 25.5 (3.1 to 47.9) | 61.1 (34.9 to 87.3) | 4.2 |
| 75 - 79 | 76.6 (19.9 to 133.4) | 7.1 (-6.8 to 21.1) | 34.5 (10.6 to 58.4) | 10.8 |
| 80 & over | 77.7 (20.1 to 135.3) | 0.0 | 31.1 (8.1 to 54.2) | - |
| Total | 34.6 (30.5 to 38.7) | 7.8 (5.9 to 9.7) | 20.7 (18.5 to 22.9) | 4.4 |

\*Suicide incidences were calculated based on the suicides occurring in 2012

### Lifetime Recalled Repetition Study

Life time previous DSH was recorded in 433 (male 47% and female 53%) randomly selected cases. Forty one (9.5%) had life time history of DSH attempts; 20 (48.8%) males and 21 (51.2%) females. The average age of cases who had made previous attempts was 26.9 years (SD 13.1, 95% CI 22.8 - 31.1). Of amongst the cases who had made previous attempts, a majority (32, 78%) had made only one previous attempt. Eight (19.5%) had two previous attempts and one (2.3%) had four.

## Discussion

The population based DSP incidence reported in our study area of 248.3/100,000 is considerably lower than that observed by Knipe et. al. (10). This difference is most likely due to double counting of DSP due to high rates of inter-hospital transfer(11)that artificially in inflates the incidence. When transfers were included and double counted DSP incidence increases to 347.4/100,000 (95% CI 338.3 – 356.4) in KD. In contrast to suicide incidence, DSP incidence among males was slightly higher than females, and the male to female ratio was 1.1:1 which, is exact opposite of the sex ratio of the district’s population (12). This finding is compatible with the previous findings, that sixteen out of seventeen studies reported higher male to female gender ratio for DSH (13). The pattern observed for age standardized DSP incidents is different to the pattern of suicide. One third of DSP occurred in 15-24 years age group, more than half less than 34 years. That follows national pattern (14) as well as pattern in South East Asian region (15, 16). This age, gender pattern can be partially explained by main six culture specific factors; (1) adolescents are often faced with stressful academic and familial expectations despite limited resources and opportunities (17) (2) socio-cultural concept related to the response towards suicidal behaviour based on sympathetic grounds (18), (3) it is a response to stressful events, which carries a powerful message to a specific person or to the society, or simply a way to conveying misgiving, anger, sadness, hopelessness, frustration, especially among adolescents(19), (4) a learned way of manipulating a situation to their own advantage, or communicating distress(19), (5) the blend of socio-economic demands and substance/alcohol misuse behaviour that associate with traditional male gender, especially after adolescence/marriage(20), and, (6) Societal attitude towards female adolescents and familial restrictions on behaviours of adolescent girls (21).

The findings of the present study indicate that self-reported recalled life-time repetition rate is 9.5%(95% CI 6.7-12.2%). Examination of records confirmed that repetition rates at four weeks, one year and two year were 1.9, 5.6 and 7.9 per hundred patients in KD, respectively. The life time repetition rate is higher compared to one year or two year repetition rate because repetitive attempts can occur at any point of life (22-24). Another potential reason for this is that the life time repetition rate was based on a referred hospital sample, which may have introduced a referral bias for patients with higher intent, whereas other rates were calculated for the entire KD, including patients presenting to primary rural hospitals many of who were not transferred to referral hospitals.

Almost all the previous studies conducted in SL were reported self-reported, life-time, recalled repetition rates. Though the method is different in this study, the self-reported life-time repetition rate of KD is close to the value reported from a socio-economically similar agricultural area published in a previous study, of 8.7% in North-Central Province (NCP)(25) and 7% in the Central and North Western Provinces(26). Two psychological autopsy studies conducted in the NCP (27) and Rathnapura (28) reported a higher lifetime value, of 26%. A telephone interview based study conducted at T.H. Peradeniya, SL, reported recalled one year repetition rate, 2.7% (29). Contextual and methodological difference partially explains the difference in rates.

Literature shows that repetition rate and excess risk carried by the previous attempt in a community differ on culture, geographical, location, outcome of suicidal behavior and the period of follow-up. A recent meta-analysis reported that, the estimated one year non-fatal repeat self-harm rate was considerably lower in Asian countries than in Western countries, 10% vs 16.3% (4). Similarly, lower repetition frequencies were reported among non-Western immigrants by a study conducted in seven European countries (30). Most of the Western countries reported higher repetition rates despite of their well developed medical, psychological and social services, compared to Asian countries including SL. It is possible that this is due to better ascertainment of cases gained through utilizing better medical records. However, with the robust methodology, our study’s results reconfirms lower rates of repetition in Asia with considerable accuracy. Rates of repetition resulting in deaths are comparable in our study with those seen in the west.

Observed low rates of repetition may be partially explained through the synergistic effect of four main culture specific pillars; (1) characteristics of risk factors associated with suicidal behavior, (2) experiences faced at the initial post attempt period, (3) effect of attempt as a solution to the trigger, and, (4) continuing extended family support. However, in depth qualitative analysis is necessary to explain these cultural factors that are responsible for lower repetition rates compared to west.

1. Compared to the Western countries, involvement of risk factors in suicidal behavior may be different in Asian countries, including Sri Lanka(31). However, exact factors and their effect on lower repetition rate should be further explored.
2. Experiences faced at the initial post attempt period may have a robust effect on reducing repetitive attempts. A study reported that nearly half lost the wish to die after surviving the act (Hettiarachchi & Kodituwakku, 1989). A majority of the attempts take place at or around the victim’s premises (32). Therefore, the situation is handled by the family members, relatives or other close individuals, up to the hospital admission. Moreover, the acts might improve the cohesion within the family at least for a short period and thus may prevent future events (33, 34). A significant proportion of all categories of health personnel expressed non-sympathy towards DSH patients (26). This non-sympathetic attitude may cause reluctance to seek health care. A significant proportion of those with DSH did not intend to end the life, but to change the situation on their advantage; therefore, they expect to seek medical interventions following attempts. In KD, 56% of patients thought that death would be unlikely if he/she received medical attention (5). Hence, the non-sympathetic attitude may discourage repetitive attempts. Though the repetitive attempts are lesser, repetitive suicidal ideation and threats may not be less.
3. A study conducted in Southern Sri Lanka revealed that, both boys and girls described suicidal attempt as a ‘quick fix’ to difficult interpersonal circumstances and visualized positive outcomes of it (35). Sri Lankan socio-cultural concept related to the response towards suicidal behavior based on sympathetic grounds. And, 26% believed in solving the problem, arranging a marriage and fulfilling wishes, as the appropriate response (18). And, parents may change their parenting strategies to more supportive parenting strategies(36). Removal of the triggering factor may prevents repetitive suicidal behavior at least for a certain period.
4. Continuing extended family support may be a factor that helps to keep lifetime repetition rate at a lower level, which has been described as a potent psycho-therapeutic factor in Indian context (37, 38). Sri Lankan socio-cultural concept related to the response towards suicidal behavior based on sympathetic grounds. Examination of culture, gender and suicidal behaviour in Sri Lanka has suggested that both emotion focused and problem focused support is deemed needed for people who have attempted suicide, with a greater emphasis on emotion-focused support for females (18). Continuing family support throughout the adolescent years and after marriage through the extended family is an integral part of the Sri Lankan culture. Further, significant proportion give higher priority to family’s requirements compared to their own needs. Therefore, majority of school adolescents perceived their families as intimate and close (60 %) and considered family as refuge (52%) for a problem (39). This may ensure the emotional warmth and bonds among the family member. Social support is well known protective factor for suicidal behavior. Hence, it might contributes to lower repetition rates.

A majority of the victims who had repetitive attempts were males, 56%. The opposite pattern was reported in NCP, female 61% (25). However, some authors reported that there was no significant difference across genders (40). Relatively higher repetition rates were reported among younger age groups in females and opposite pattern in males. Literature on psychological and socio-economical predictors of repetition showed that they are not different from the risk-factors of non-repetitive self-harm behavior (41).

The risk of repetition is higher in initial post event period. The median times to repetition within 1 year and 2 years were, 92 (IQR 10 - 238) and 191 (IQR 29 - 419.5) days respectively. The risk for repetition is highest in the first 3 to 6 months after a suicide attempt, but remained substantially elevated from the general population for at least 2 years (Bridge et al., 2006; Goldston et al., 1999; Lewinsohn, Rohde, & Seeley, 1996). The median time to repetition within 1 year was 105 days in Taiwan (Kwok, Yip, Gunnell, Kuo, & Chen, 2014).

The longer lengths of hospital stay in SL hospitals have been proposed by authors as a reason for observed difference with western countries (25). Similar to the current study findings, previous work has shown that the initial post attempt period carries the highest risk for repetition (42). Short hospital stay may expose the patient to the same environment that lead to the suicidal behavior. In England, half of the self-harm patients presenting to the emergency department were discharged without being admitted to hospital (43). In contrast, same day discharges were limited to 3% and 4% in THK and peripheral hospitals. The median hospital stay was reported as one day in western countries (4, 44, 45), whereas it was 2 days in the KD. Hence, longer length of hospital stay can be considered as a valid argument for a lower repetition rate compared to western countries.

Higher case fatality of the first episode was a suggested explanation for the observed lower repetition rate (25). This explanation seems unlikely as there has been a significant reduction in case fatality rates over time whereas repetition rates have remained low. (10). If we assume all 221 suicides with poisoning, reported to police stations in 2011, survived following the initial event and had a repetitive event within a year, there would be 448 repetition cases within a year. Then, the one year repetition rate would be 11.1%, 95% CI: 10.2% to 12.1% and the value much lower to the one year repetition rate in Europe.

Suicide incidence of KD has been remained stable over the last decade, in 2012,20.7/100,000 and the average value for 2001-2006 period, 21.3/100,000(14, 46) in the presence of rising trends of DSP(10). This observation can be explained by three main mechanisms; (1) continuum of the reduction observed from 1996 with some of the actions taken in 1990s: such as restricting the import and sale of WHO Class I toxicity pesticides and decriminalization of suicide, (2) improvements of self-poisoning medical management, and, (3) shifting of methods from lethal pesticides to less lethal medicaments(10, 47). Male to female ratio of suicide incidences is similar to Europe and countries in American subcontinent, rather than Asian countries(15).Age standardized suicide incidence pattern is similar to national (14) as well as patter in most parts of the world (48-52).

Though there is a considerable amount of literature available to explain the risk factors that are responsible for higher rates of DSH in Sri Lanka, culture specific protective factors that leads to lower repetition rates are poorly explored. These protective factors should be further explored to explain the lower rates of repetition. These protective factors may provide a base to promising preventive strategies of DSH. Further, measures directed to prevention of repetition alone may not produce considerable impact on preventing suicidal behavior, in the presence of lower repetition rates.

### Limitations

Data collection was conducted only in government hospitals. Less severe cases those may present to private-sector out-patient-care services might missed from data collection.

In calculating record based one and two year repetition rates; only the DSP admissions were considered for non-fatal events; not considering other methods of DSH, might have had an effect on lower rates. It has been suggested that individuals who attempt self-injury are more prone to repetitive attempts compared to those who attempt self-poisoning (53). However, this effect may not be be significant because more than 80% of DSH are due to poisoning and over dosage of medicines, and all fatal events were considered.

There are no unique patient identifiers in the provincial or national health system in Sri Lanka. The source of information, for the patient details, at the point of patient registration is the guardian and/or patient and there are no verifications or cross-checking. Because of the stigma, they might provide incomplete information to hide their identity. Therefore, validity and reliability of the identification details in heath records may be limited. This may have affected the reliability of repetition matching process.

## Conclusion

Repetition rate of DSH in Sri Lanka is very low, compared to Western countries and other countries in the region. Therefore, measures directed to prevention of repetition alone may not produce significant impact on preventing suicidal behavior. Culture specific protective factors that leads to lower repetition rates should be further explored and they may provide a base for promising preventive strategies of DSH.

## Supplemental tables

**Table 2:**
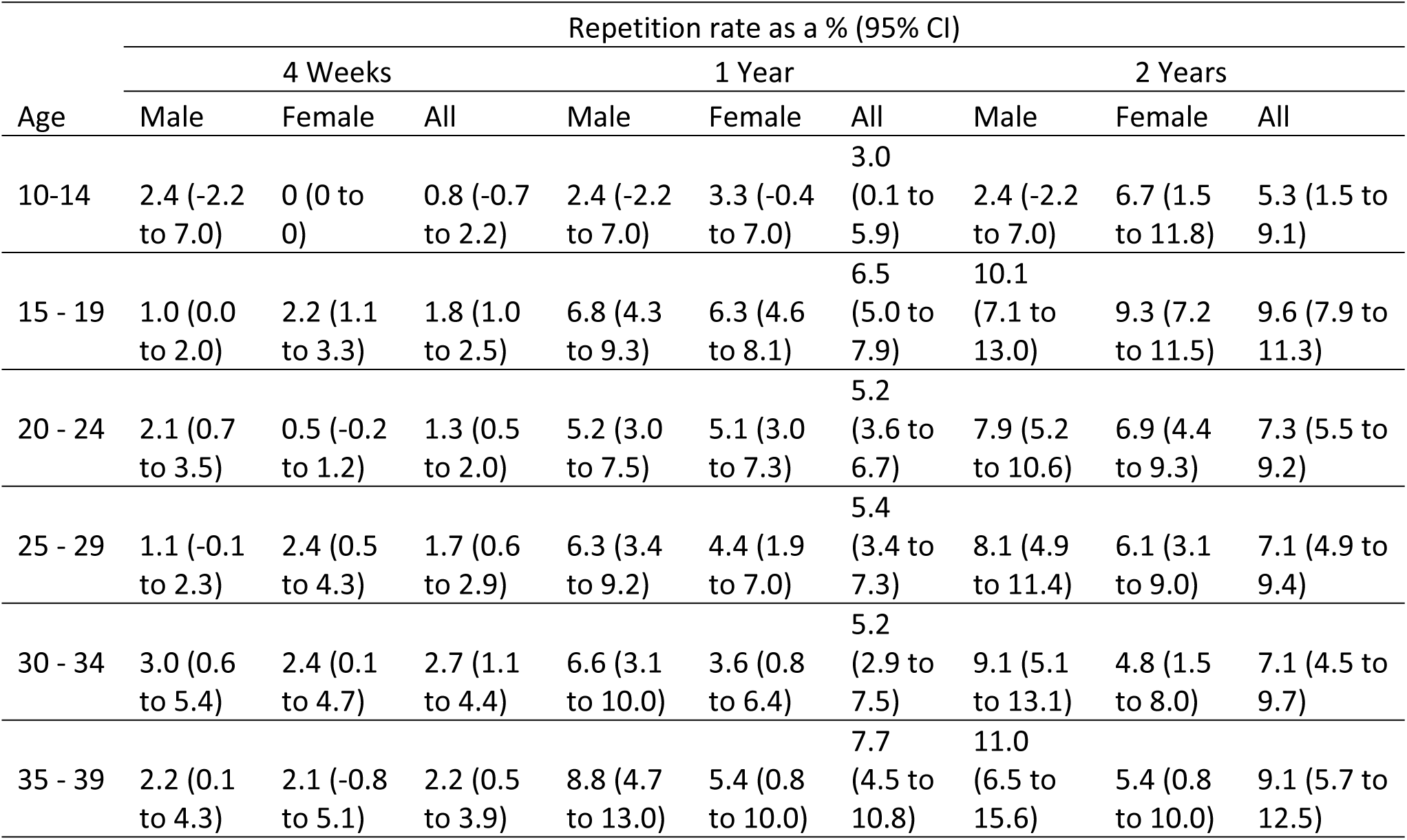
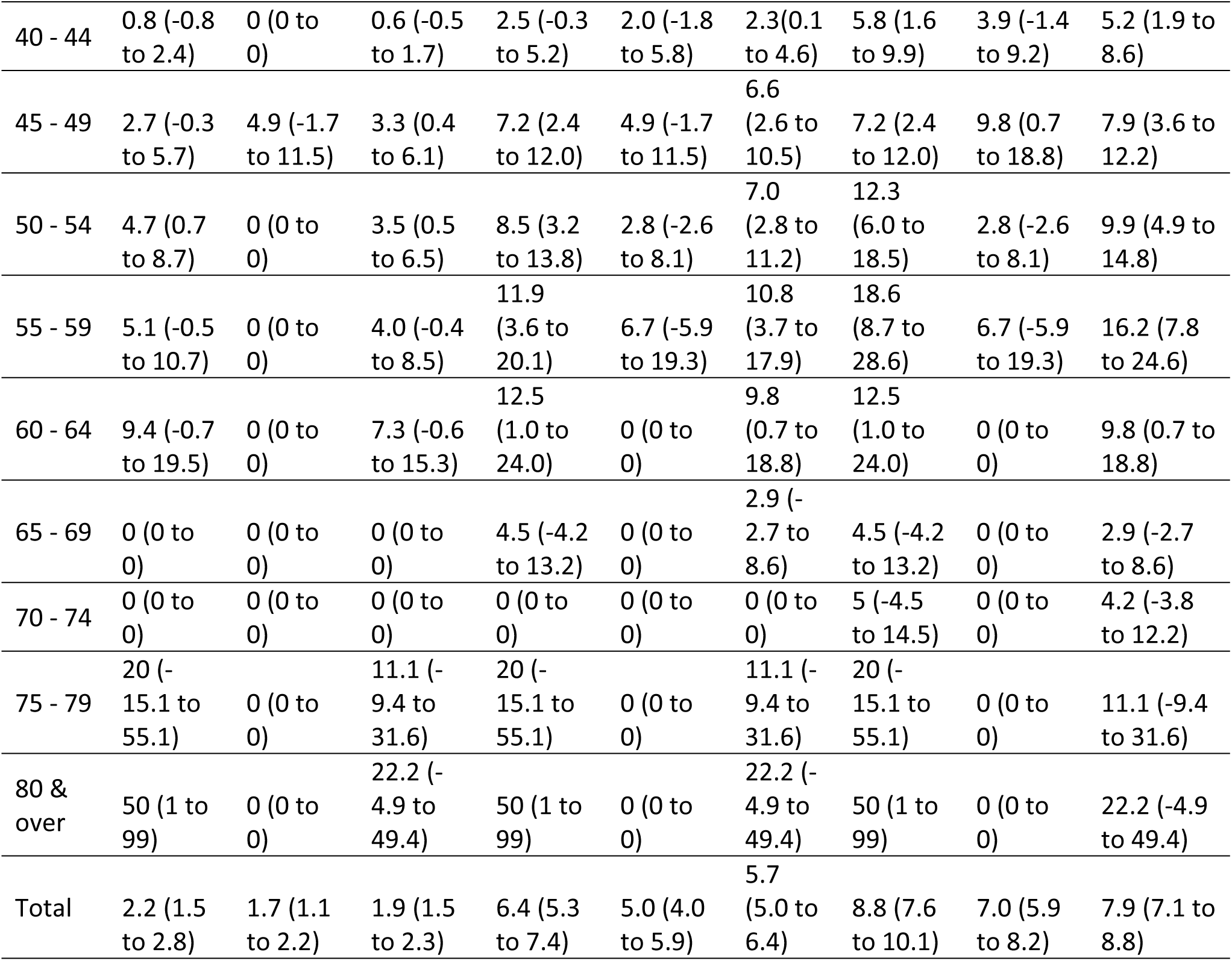
four weeks, one year and two year repetition rates by age and sex

## References

1. Bertolote JM, Fleischmann A. A global perspective on the magnitude of suicide mortality. In: Wasserman D, Wasserman C, editors. Oxford textbook of suicidology and suicide prevention: a global perspective. illustrated ed: Oxford University Press; 2009. p. 91–8.

2. Bertolote JM, Fleischmann A. A global perspective in the epidemiology of suicide. Suicidologi. 2002;7(2):7–9.

3. La Vecchia C, Lucchini F, Levi F. Worldwide trends in suicide mortality, 1955-1989. Acta Psychiatrica Scandinavica. 1994;90(1):53–64.

4. Carroll R, Metcalfe C, Gunnell D. Hospital Presenting Self-Harm and Risk of Fatal and Non-Fatal Repetition: Systematic Review and Meta-Analysis. PLoS ONE. 2014;9(2):e89944.

5. Pushpakumara PHGJ. Epidemiological Pattern And Associates of Deliberate Self-Harm in Kurunegala District, Sri Lanka. Peradeniya, Sri Lanka: Faculty of Medicine, University of Peradeniya; 2017.

6. Department of Census and Statistics Sri Lanka. Statistical Handbook: Kurunegala District - Population by Age groups and Gender within District - 2014 (Table: 2 .8). Colombo, Sri Lanka 2015 [cited 2016]; Available from: http://www.statistics.gov.lk/statistical%20Hbook/2015/Kurunegala/Table%202.8.pdf.

7. Department of Census and Statistics Sri Lanka. Statistical Handbook: Kurunegala District - Information on Government Hospitals by Divisional Level - 2014 (Table: 8.1). Colombo, Sri Lanka: Department of Census and Statistics Sri Lanka; 2015; Available from: http://www.statistics.gov.lk/statistical%20Hbook/2015/Kurunegala/Table%208.1.pdf.

8. Hettige B, Karunananda AS, editors. Transliteration System for English to Sinhala Machine Translation. Second International Conference on Industrial and Information Systems (ICIIS 2007); 2008 8-11 August 2007; University of Peradeniya, Sri Lanka.

9. Fernando SC, Dias G, editors. Inexact Matching of Proper Names in Sinhala. IEEE Reglon 10 Colloquium and Th’ird Internatlonal Conference on Industrial and Informat’ion Systems (ICIIS-2008); 2007 December 8-10, 2008; Indian Institute of Technology, Kharagpur, India.

10. Knipe DW, Metcalfe C, Fernando R, Pearson M, Konradsen F, Eddleston M, et al. Suicide in Sri Lanka 1975–2012: age, period and cohort analysis of police and hospital data. BMC public health. 2014;14(1):839.

11. Senarathna L, Buckley N, Jayamanna S, Kelly P, Dibley M, Dawson A. Validity of referral hospitals for the toxicovigilance of acute poisoning in Sri Lanka. Bull World Health Organ. 2012;90(6):436–43A.

12. Department of Census and Statistics Sri Lanka. Chapter-2: Information related to Population Housing in District Statistical Hand Book Kurunegala. Colombo: Department of Census and Statistics-Sri Lanka, 2013.

13. Rajapakse T, Griffiths KM, Christensen H. Characteristics of non-fatal self-poisoning in Sri Lanka: a systematic review. BMC public health. 2013;13:133.

14. Sri Lanka Police. Sri Lanka Police: Crime Trends (Year 2005 to 2015). Colombo: Information Technology Division, Sri Lanka Police; 2016 [updated 19.3.2016; cited 2016 20.3.2016]; Available from: http://www.police.lk/index.php/crime-trends.

15. Varnik P. Suicide in the world. International journal of environmental research and public health. 2012;9(3):760–71. Epub 2012/06/13.

16. Khan MM. Suicide on the Indian Subcontinent. Crisis: The Journal of Crisis Intervention and Suicide Prevention. 2002;23(3):104–7.

17. Nastasi BK, Hitchcock JH, Burkholder G, Varjas K, Sarkar S, Jayasena A. Assessing Adolescents’ Understanding of and Reactions to Stress in Different Cultures: Results of a Mixed-Methods Approach. School Psychology International. 2007;28(2):163–78.

18. Marecek J. Culture, gender, and suicidal behavior in Sri Lanka. Suicide and Life-Threatening Behavior. 1998;28(1):69–81.

19. Konradsen F, Hoek Wvd, Peiris P. Reaching for the bottle of pesticide—A cry for help. Self-inflicted poisonings in Sri Lanka. Social Science & Medicine. 2006;62(7):1710–9.

20. Marecek J. Young Women’s Suicide In Sri Lanka: Cultural, Ecological, And Psychological Factors. Asian Journal of Counselling. 2006;13(1):63–92.

21. Houle J, Mishara BL, Chagnon F. An empirical test of a mediation model of the impact of the traditional male gender role on suicidal behavior in men. Journal of Affective Disorders. 2008;107(1-3):37–43.

22. Bridge JA, Goldstein TR, Brent DA. Adolescent suicide and suicidal behavior. Journal of Child Psychology and Psychiatry. 2006;47(3-4):372–94.

23. Goldston DB, Daniel SS, Reboussin DM, Reboussin BA, Frazier PH, Kelley AE. Suicide attempts among formerly hospitalized adolescents: A prospective naturalistic study of risk during the first 5 years after discharge. Journal of the American Academy of Child and Adolescent Psychiatry. 1999;38:660–71.

24. Lewinsohn PM, Rohde P, Seeley JR. Adolescent suicidal ideation and attempts: Prevalence, risk factors, and clinical implications. Clinical Psychology Science and Practice. 1996;3(25–36).

25. Mohamed F, Perera A, Wijayaweera K, Kularatne K, Jayamanne S, Eddleston M, et al. The prevalence of previous self-harm amongst self-poisoning patients in Sri Lanka. Social psychiatry and psychiatric epidemiology. 2011;46(6):517–20. Epub 2010/04/08.

26. de Silva HJ, Kasturiarachchi N, Seneviratne SL, Senaratne DC, Molagoda A, Ellawala NS. Suicide in Sri Lanka: points to ponder. Ceylon Medical Journal. 2000;45:17–24.

27. Abeyasinghe R, Gunnell D. Psychological autopsy study of suicide in three rural and semi-rural districts of Sri Lanka. Social psychiatry and psychiatric epidemiology. 2008;43(4):280–5. Epub 2008/02/07.

28. Samaraweera S, Sumathipala A, Siribaddana S, Sivayogan S, Bhugra D. Completed suicide among Sinhalese in Sri Lanka: a psychological autopsy study. Suicide & life-threatening behavior. 2008;38(2):221–8.

29. Rajapakse TN, Griffiths KM, Cotton S, Christensen H. Repetition rate after non-fatal self-poisoning in Sri-Lanka: a one year prospective longitudinal study. The Ceylon Medical Journal. 2016(61):154–8.

30. Lipsicas CB, Mäkinen IH, Wasserman D, Apter A, Kerkhof A, Michel K, et al. Repetition of Attempted Suicide Among Immigrants in Europe. Can J Psychiatry. 2014;59(10):539–47.

31. Chen Y-Y, Chien-Chang Wu K, Yousuf S, Yip PSF. Suicide in Asia: Opportunities and Challenges. Epidemiologic reviews. 2012;34(1):129–44.

32. Mohamed F, Manuweera G, Gunnell D, Azher S, Eddleston M, Dawson A, et al. Pattern of pesticide storage before pesticide self-poisoning in rural Sri Lanka. BMC public health. 2009;9:405. Epub 2009/11/06.

33. Franic T, Dodig G, Kardum G, Marcinko D, Ujevid A, Bilušid M. Early Adolescence and Suicidal Ideations in Croatia: Sociodemographic, Behavioral, and Psychometric Correlates. Crisis: The Journal of Crisis Intervention and Suicide Prevention. 2011;32(6):334–45.

34. Lee AY, Pridmore S. Suicide and gender ratios in Tasmania (Australia) using the Operationalized Predicaments of Suicide tool, and negative experiences. Australasian Psychiatry. 2014;22(2):140–3.

35. Senadheera C, editor. Deliberate self-harm of children and adolescents: a hospital based study. Suicide in Sri Lanka: Past, Present and Future Transformations; 2013; University of Colombo: Department of Sociology, University of Colombo.

36. Ferrey AE, Hughes ND, Simkin S, Locock L, Stewart A, Kapur N, et al. Changes in parenting strategies after a young person’s self-harm: a qualitative study. Child Adolesc Psychiatry Ment Health. 2016;10(20).

37. Sethi B. FAMILY AS A POTENT THERAPEUTIC FORCE. Indian Journal of Psychiatry. 1989;31(1):22–30.

38. Chadda RK, Deb KS. Indian family systems, collectivistic society and psychotherapy. Indian Journal of Psychiatry. 2013;55(Suppl 2):S299–309.

39. Thalagala N, Rajapakse L, Yakandawala H. National Survey on Emerging Issues among Adolescents in Sri Lanka. Colombo, Sri Lanka: UNICEF Sri Lanka, 2004.

40. Nordentoft M, Branner J. Gender differences in suicidal intent and choice of method among suicide attempters. Crisis: The Journal of Crisis Intervention and Suicide Prevention. 2008;29(4):209–12.

41. Larkin C, Blasi ZD, Arensman E. Risk Factors for Repetition of Self-Harm: A Systematic Review of Prospective Hospital-Based Studies. PLoS ONE. 2014;9(1):e84282.

42. Gilbody S, House A, Owens D. The early repetition of deliberate self harm. J R Coll Physicians Lond. 1997;31(2):171–2.

43. Gunnell D, Bennewith O, Peters TJ, House A, Hawton K. The epidemiology and management of self-harm amongst adults in England. J Public Health (Oxf). 2005;27(1):67–73.

44. Carter GL, Whyte IM, Ball K, Carter NT, Dawson AH, Carr VJ, et al. Repetition of deliberate self-poisoning in an Australian hospital-treated population. Med J Aust. 1999;170(7):307–11.

45. Gunnell DJ, Brooks J, Peters TJ. Epidemiology and patterns of hospital use after parasuicide in the south west of England. J Epidemiol Community Health. 1996;50(1):24–9.

46. Registrar General’s Department-Sri Lanka. Bulletin of Vital Statistics. Colombo: Registrar General’s Department, Ministry of Public Administration & Home Affairs-Sri Lanka, Department RGs; 2010.

47. de Silva VA, Senanayake SM, Dias P, Hanwella R. From pesticides to medicinal drugs: time series analyses of methods of self-harm in Sri Lanka. Bulletin of the World Health Organization. 2012;90:40–6.

48. Vijayakumar L. Suicide in Asia: Causes and Prevention. Hong Kong, China: Hong Kong University Press; 2008.

49. Hawton K, Van Heeringen K. Suicide. Lancet. 2009;373(9672):1372–81.

50. De Leo D, Padoani W, Scocco P, Lie D, Bille-Brahe U, Arensman E, et al. Attempted and completed suicide in older subjects: results from the WHO/EURO Multicentre Study of Suicidal Behaviour. International Journal of Geriatric Psychiatry. 2001;16(3):300–10.

51. Dombrovski AY, Szanto K, Duberstein P, Conner KR, Houck PR, Conwell Y. Sex Differences in Correlates of Suicide Attempt Lethality in Late Life. American Journal of Geriatric Psychiatry. 2008;16(11):905–13.

52. Wasserman D, Cheng Q, Jiang G-X. Global suicide rates among young people aged 15-19. World Psychiatry. 2005;4:114–20.

53. Lilley R, Owens D, Horrocks J, House A, Noble R, Bergen H, et al. Hospital care and repetition following self-harm: multicentre comparison of self-poisoning and self-injury. The British Journal of Psychiatry. 2008;192(6):440–5.

